# Development of a quantitative PCR assay for the detection and enumeration of a potentially ciguatoxin-producing dinoflagellate, *Gambierdiscus lapillus* (Gonyaulacales, Dinophyceae)

**DOI:** 10.1101/544247

**Authors:** A.L. Kretzschmar, A. Verma, G.S. Kohli, S.A. Murray

**Affiliations:** Climate Change Cluster (C3), University of Technology Sydney, Ultimo, 2007 NSW, Australia; ithree institute (i3), University of Technology Sydney, Ultimo, 2007 NSW, Australia; Alfred Wegener-Institut Helmholtz-Zentrum fr Polar- und Meeresforschung, Am Handelshafen 12, 27570, Bremerhaven, Germany

**Keywords:** Ciguatera fish poisoning, *Gambierdiscus lapillus*, Quantitative PCR assay, Great Barrier Reef

## Abstract

Ciguatera fish poisoning is an illness contracted through the ingestion of seafood containing ciguatoxins. It is prevalent in tropical regions worldwide, including in Australia. Ciguatoxins are produced by some species of *Gambierdiscus.* Therefore, screening of *Gambierdiscus* species identification through quantitative PCR (qPCR), along with the determination of species toxicity, can be useful in monitoring potential ciguatera risk in these regions. In Australia, the identity, distribution and abundance of ciguatoxin producing *Gambierdiscus* spp. is largely unknown. In this study we developed a rapid qPCR assay to quantify the presence and abundance of *Gambierdiscus lapillus*, a likely ciguatoxic species. We assessed the specificity and efficiency of the qPCR assay. The assay was tested on 25 environmental samples from the Heron Island reef in the southern Great Barrier Reef, a ciguatera endemic region, in triplicate to determine the presence and patchiness of these species across samples from *Chnoospora* sp., *Padina* sp. and *Sargassum* macroalgal hosts.

**Author’s summary:** Ciguatera fish poisoning is a human disease contracted by ingesting seafood contaminated with a group of neurotoxins. The group of neurotoxins, named ciguatoxins, are synthesised by species of single celled marine algae from the genus *Gambierdiscus.*

Ciguatera fish poisoning occurs worldwide, particularly in tropical nations. Pacific Island nations are disproportionately impacted, and this impact is predicted to increase as the effects of climate change unfold. Few effective monitoring and mitigation strategies exist for ciguatera fish poisoning, and reporting rates of the disease are estimated to be approximately 20% at best. A global ciguatera strategy was developed by a group of researchers coordinated by UNESCO’s Intergovernmental Oceanographic Commission to characterise the cause and mode of action of ciguatera fish poisoning, as a matter of urgency.

In this study, we designed a qPCR assay to detect a species of microalgae, *Gambierdiscus lapillus. Gambierdiscus lapillus* produces compounds with ciguatoxin-like properties, which may lead to ciguatoxin uptake in fish in the Australian region. This assay was sensitive and able to detect the presence of *Gambierdiscus lapillus* in a range of environmental samples from the Great Barrier Reef region, Australia.

## Introduction

Benthic dinoflagellates of the genus *Gambierdiscus* Adachi & Fukuyo produce ciguatoxins (CTX), which can accumulate in humans via consumption of contaminated seafood and cause ciguatera fish poisoning (CFP) (Fig. 1).

**Figure 1:**
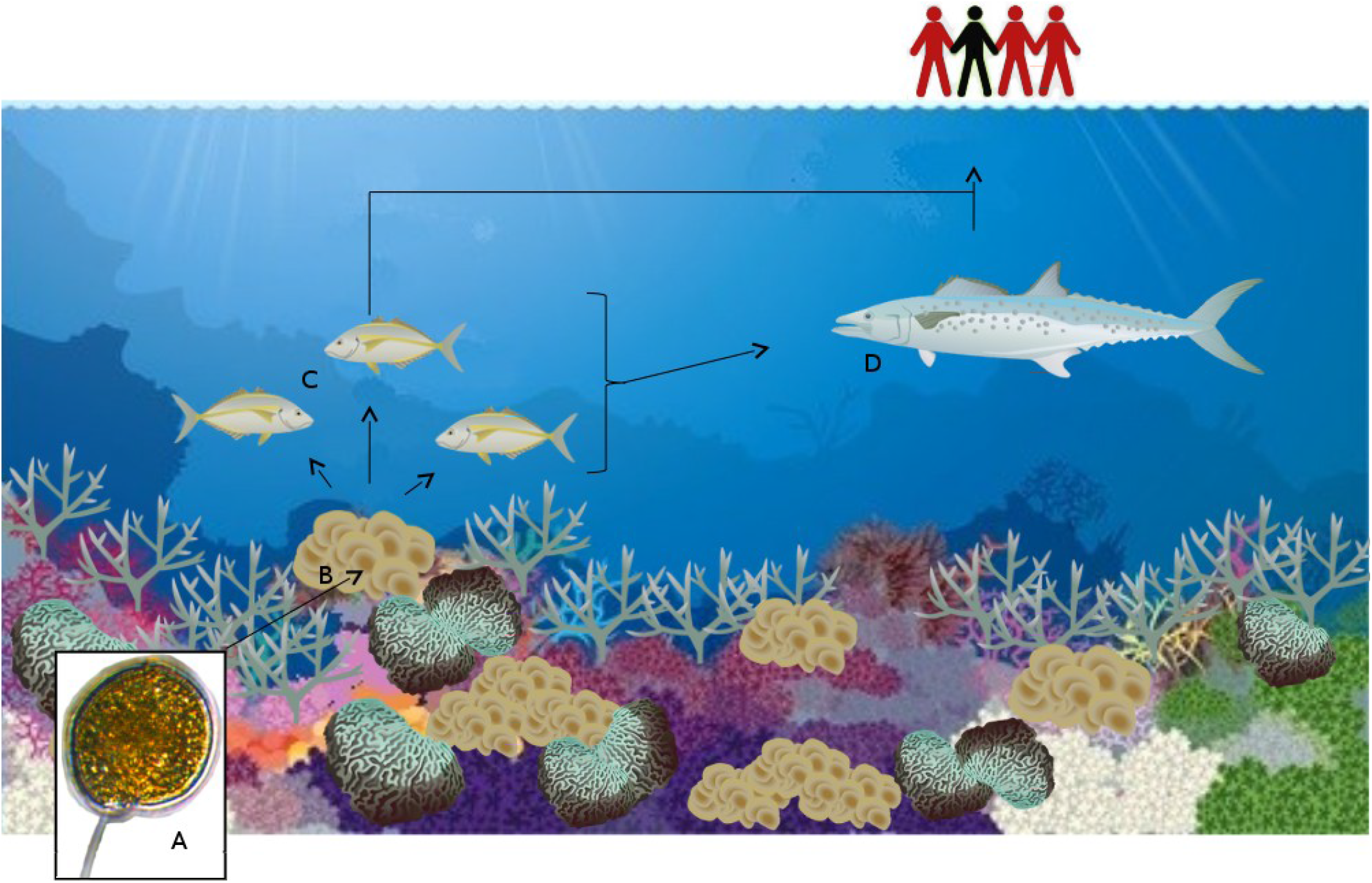
The mechanism of bioaccumulation of CTXs, with *Gambierdiscus* (for example G. *polynesiensis* (A)) at the base of the food web inhabiting the macroalgae *Padina* spp. (B) [1]. A herbivore, here a white trevally *(Pseudocaranx dentex)* (C) [2] consumes CTX from G. *polynesiensis* along with the macroalgae, which is then either passes directly to humans through consumption, or through an intermediary piscivorous vector such as Australian spotted mackerel *(Scomberomorus munroi)* (D) [3]. Image of G. *polynesiensis* (strain CG15) taken by A. L. Kretzschmar, 2016, Nikon Eclipse TS100 equipped with an Infinite Luminera 1 camera.

The symptoms of CFP are largely gastrointestinal and neurotoxic, however, in severe cases, further complications such as cardiovascular or severe neurological symptoms can appear [4]. Species of *Gambierdiscus* spp. are predominantly epiphytic, growing on macroalgae and other substrates such as coral detritus. Species of *Gambierdiscus* spp. can vary in the production of CTXs and/or maitotoxins (MTXs) [5, 6]. If a particular *Gambierdiscus* sp. is a CTX producer, and inhabit a palatable macroalgal substrate, the toxins bioaccumulate in herbivorous fish and filter feeders with the potential to travel up the food chain to cause CFP in humans [7, 8].

*Gambierdiscus* was first identified in 1977, with the type species G. *toxicus* Adachi & Fukuyo [9]. The genus remained monotypic for 18 years until the discovery of a second species G. *belizeanus* Faust [10]. To date, the genus comprises 14 described species and 6 ribo/species types [9, 10, 11, 12, 13, 14, 15, 16, 17, 18, 19, 20, 21, 22]. A major revision of the *Gambierdiscus* species taxonomy was undertaken by Litaker et al. (2009). Reports of *Gambierdiscus* spp. identified based on morphology alone, prior to this revision; need to be considered with caution as several new *Gambierdiscus* spp. were defined [23, 24, 25]. Further, intra-species variation and inter-species similarities can cause misidentification [6, 19, 26]. Hence, molecular genetic tools are important for determining the distribution and abundance of *Gambierdiscus* species and assess the risk of CFP in that region [6, 19].

*Gambierdiscus* spp. produce a suite of different polyketide compounds – CTX, maitotoxin (MTX), gambierone, gambieric acid and gambierol have been characterised to date [27, 28, 29, 30, 31]. While any of these can contribute to toxicity, only CTX has been clearly linked to CFP in humans [7, 8]. The toxin profile of many *Gambierdiscus* species is not well understood, and many different assays have been used to determine CTX toxicity [32]. Assays, such as mouse bioassays and neuroblastoma cell-line bioassays are good indicators of the toxicity of an organism, however species/strain specific toxin profiles needs to be elucidated with LC-MS/MS in order to characterise individual toxin congeners [33]. The toxin profile of *Gambierdiscus polynesiensis* Chinain & Faust is one of the only *Gambierdiscus* spp. whose production of CTX congeners (P-CTX-3B, P-CTX-3C, P-CTX-4A, P-CTX-4B and M-seco-CTX-3C) has been verified by LC-MS/MS in isolates from French Polynesia and the Cook Islands, and is thought to be the principal cause of CFP in the Pacific region [34, 35]. However recently, a G. *polynesiensis* strain isolated from the Kermandec Islands, Pacific Ocean, did not exhibit CTX toxicity detectable by LC-MS/MS [36]. An uncharacterised peak in the CTX phase of several strains of *Gambierdiscus lapillus* extracts was reported via LC-MS/MS, which did not match any available CTX standards (CTX-3B, CTX-3C, CTX-4A, CTX-4B) [19]. Further, Larsson et al. (2018) found that G. *lapillus* extracts showed CTX-like activity when investigated with a bioassay. Therefore, this species likely produces previously uncharacterised CTX congener(s), and its production of CTX compounds requires further investigation. Determining the toxin profile of *Gambierdiscus* species requires toxin standards for comparative peak analysis. However, these are currently not commercially available. Therefore, progress in determining the toxins produced by species of *Gambierdiscus* has been comparatively slow, though bioassays provide a strong indicator for toxin production.

CFP was put forward as a ‘‘neglected tropical disease” by expert researchers in this area, supported by the Intergovernmental Oceanographic Commissions (IOC) Intergovernmental Panel on Harmful Algal Blooms (IPHAB), as part of the United Nations Educational, Scientific and Cultural Organization), and a global ciguatera strategy was developed [32]. One element of the IOC/IPHAB Global Ciguatera Strategy is to investigate various species of the genus *Gambierdiscus*, determine which species produce CTXs through LC-MS/MS and other means, and develop efficient and reliable molecular monitoring tools for the species of interest [32]. Quantitative PCR (qPCR) is a useful molecular genetic screening tool, as it can give species-specific and quantitative results from DNA samples extracted from environmental samples [32].

Currently there is one qPCR assay to identify the presence of the genera *Gambierdiscus/Fukuyoa* [37]. Assays for species specific identification are available for 9 of the 14 described *Gambierdiscus* spp. and 3 out of 6 undescribed *Gambierdiscus* sp. types/ribotypes (Table 1). It is noteworthy that the qPCR assays described by Darius et al. (2017) rely on species identification based on the melt curve of the qPCR product, which requires any subsequent users of these assays to have a reference culture for positive identification rather than rely on a positive result being linked to the species investigated. Assays are available for 2 of the 3 species of *Fukuyoa* (Table 1), which seceded from *Gambierdiscus* as their own genus in 2015 [38]. *Fukoyoa* spp. are of interest for monitoring purposes as they produce MTXs, however the involvement of MTXs in CFP has not been resolved yet [39].

**Table 1:** Published qPCR assays for *Gambierdiscus* and *Fukoyoa* spp.

| Species | Method | Reference |
| --- | --- | --- |
| ***Gambierdiscus* spp.** |  |  |
| *G. australes* | TaqMan Probes & SYBR Green | [40, 41] |
| *G. belizeanus* | SYBR Green | [42] |
| *G. caribaeus* | SYBR Green | [42] |
| *G. carolinianus* | SYBR Green | [42] |
| *G. carpenteri* | SYBR Green | [42] |
| *G. pacificus* | SYBR Green | [41] |
| *G. polynesiensis* | SYBR Green | [41] |
| *G. scabrosus* | TaqMan Probes | [40] |
| *G. toxicus* | SYBR Green | [41] |
| *Gambierdiscus* sp. ribotype 2 | SYBR Green | [42] |
| *Gambierdiscus* sp. type 2 | TaqMan Probes | [40] |
| *Gambierdiscus* sp. type 3 | TaqMan Probes | [40] |
| ***Fukuyoa* spp.** |  |  |
| *Fukuyoa ruetzleri* | SYBR Green | [42] |
| *Fukuyoa* cf. *yasumotoi* | TaqMan Probes | [40] |

In Australia, outbreaks of CFP occur annually in Queensland [43]. However, due to the complicated presentation of symptoms, the reporting rate is less than 20% [44]. Annually, there have been 7-69 reported cases between 2011 and 2015 (considering the report rate, > 35-345 cases, see Table 2), with 2 fatalities reported in the state [45]. Cases of CFP from Spanish Mackerel *(Scomberomorus commerson)* caught in NSW have been reported since 2014 [46], with five separate outbreaks affecting a total of 24 people since then [47]. Farrell et al. (2017) put forward a series if recommendations managing the emerging CFP risk in NSW.

**Table 2:** Cases of CFP reported to health authorities in Queensland, Australia, between 2011 and 2015, by Queensland Health [43].

| Year | 2011 | 2012 | 2013 | 2014 | 2015 |
| --- | --- | --- | --- | --- | --- |
| Recorded CFP cases | 18 | 7 | 25 | 69 | 11 |
| Extrapolated CFP incidences | ~90 | ~35 | ~125 | ~345 | ~55 |

Despite the prevalence of CFP in Australia, the characterization of *Gambierdiscus* species present in Australia is incomplete. A species that produces known CTX toxins has not been identified from Australia as yet. Larsson et al. (2018) have identified some candidate species, two of which show some CTX-like bioactivity [48]. Over 50% of Australia’s vast coastline (total 66,000 km) is tropical or subtropical, and may be considered potential habitat for *Gambierdiscus* spp. [19]. Seven species of *Gambierdiscus* have been identified from the sub-tropical east Australian coastline namely, G. *belizeanus* [49], G. *carpenteri* [6, 50], G. *honu* (based on D8-D10 LSU sequence matching to a study by Richlen et al. [51]) [18], G. *lapillus* [19, 48], G. *toxicus* [52] and two potentially new species [48], as well as *F. yasumotoi* [49]. Using high throughput amplicon sequencing, *Gambierdiscus* was identified to the genus level in Broome, Western Australia [53], indicating that this is a coastline that should be examined further for CFP risk. qPCR primers that can be used for identification in Australia for potential monitoring purposes, have been developed for G. *belizeanus*, G. *carpenteri* and *F. yasumotoi* [40, 42].

The aim of this study was to develop and test a qPCR assay to detect G. *lapillus* that exclusively amplifies the target species without requiring the operator to have a positive control for comparison. The assay then applied to environmental samples for the detection and enumeration of G. *lapillus* at Heron Island, GBR, in a region in which CFP cases are regularly reported.

## Methods

### Clonal strains and culturing conditions

Three strains of G. *lapillus* and one strain of G. cf. *silvae* were isolated from Heron Island, Australia, as previously described [19]. Two strains of G. *polynesiensis* were isolated from Rarotonga, Cook Islands (Table 3). The cultures were maintained in 5x diluted F/2 media [24] at 27 °C, 60mol •-m^2^•-s light in 12hr light to dark cycles.

**Table 3:** List of *Gambierdiscus* clonal strains used for the qPCR assay.

| Species | Collection site | Collection date | Latitude | Longitude | Strain name |
| --- | --- | --- | --- | --- | --- |
| *G. lapillus* | Heron Island, Australia | July 2014 | 23° 4420' S | 151° 9140' E | HG4 |
|  |  |  |  |  | HG6 |
|  |  |  |  |  | HG7 |
| *G. polynesiensis* | Rarotonga, Cook Islands | November 2014 | 21° 2486' S | 159° 7286' W | CG14 |
|  |  |  |  |  | CG15 |
| *G. cf. silvae* | Heron Island, Australia | July 2014 | 23° 4420' S | 151° 9140' E | HG5 |

### DNA extraction and species specific primer design

Genomic DNA was extracted using a modified CTAB method [54]. The purity and concentration of the extracted DNA was measured using the Nanodrop (Nanodrop2000, Thermo Scientific), and the integrity of the DNA was visualised on 1% agarose gel. A unique primer set was designed for the small-subunit (SSU) rDNA region of G. *lapillus* based on sequences available in the GenBank reference database (KU558929-33). The target sequences were aligned against sequences of all other *Gambierdiscus* spp. that were available on GenBank reference database, with the MUSCLE algorithm (maximum of 8 iterations) [55] used through the Geneious software, version 8.1.7 [56]. Unique sites were determined manually (Table 4). Primers were synthesised by Integrated DNA Technologies (IA, USA). The primer set was tested systematically for secondary product formation for all 3 strains of G. *lapillus* (Table 3) via standard PCR in 25μl mixture in PCR tubes. The mixture contained 0.6 μM forward and reverse primer, 0.4 μM BSA, 2 – 20 ng DNA, 12.5 μl 2xEconoTaq (Lucigen) and 7.5 μl PCR grade water. The PCR cycling comprised of an initial 10 min step at 94 °C, followed by 30 cycles of denaturing at 94 °C for 30 sec, annealing at 60 °C for 30 sec and extension at 72 °C for 1 min, finalised with 3 minutes of extension at 72 °C. Products were visualised on a 1% agarose gel.

**Table 4:**
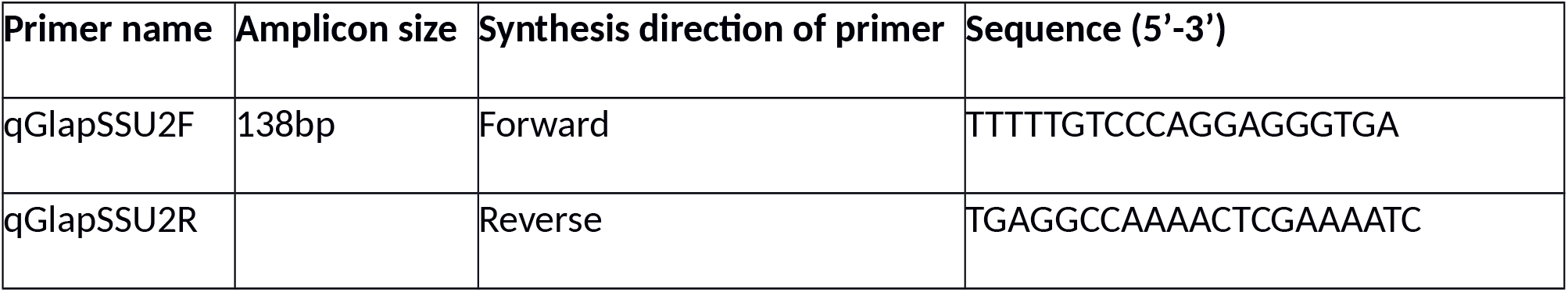
G. *lapillus* specific qPCR primer set for SSU rDNA designed in this study.

### Evaluation of primer specificity

To verify primer set specificity as listed in Table 4, DNA was extracted using CTAB buffer [57] from G. *australes* (CCMP1650 and CG61), *G. belizeanus* (CCMP401), *G. carpenteri* (UTSMER9A3), *G. pacificus* (CAWD149) and *G.* cf. *silvae* (HG5). *G. cheloniae* (CAWD232) DNA was extracted using a PowerSoil DNA isolation kit (Mo Bio Inc., CA, USA). *G. scabrosus* (KW070922_1) DNA was extracted using DNeasy Plant Mini Kit (Quiagen, Tokyo, Japan) according to the manufacturer’s protocol. For all extracted samples, the presence and integrity of genomic DNA was assessed on 1% agarose gel. The primer set designed for G. *lapillus* was tested for cross-reactivity against all other *Gambierdiscus* spp. available via PCR (BioRadT100 Thermal Cycler (CA, USA)), appropriate positive and negative controls were applied. PCR amplicons were visually assessed on 1% agarose gel.

### Evaluation of primer sensitivity

The qPCR reaction mixture contained 10 μl SYBR Select Master Mix (Thermo Fisher Scientific), 7 μl MilliQ water, 0.5 μM forward and reverse primers and 2 – 20 ng DNA template, for a final volume of 20 μl.

Cycling conditions consisted of 10 min at 95, then 40 cycles of 95 °C for 15 seconds and 60 °C for 30 seconds, followed by a temperature gradient for melt curve construction.

### Calibration curve construction

Standard curves were constructed to determine the efficiency of the assay, using a synthetic gene fragment approach, and also to use to quantify species presence, using calibration curves based on DNA extracted from known cell numbers. For curves based on synthetic gene fragments, a 10-fold serial dilution of a synthesised fragment containing the SSU target sequence, forward and reverse primer sites and 50bp flanking both primer sites matching sequencing results were generated. Cell-based standard curves were constructed using 10-fold dilutions of gDNA extract of known cell concentrations. The calibration curves for both methods were calculated (R^2^, PCR efficiency and regression line slope) and graphed in R version 3.2.3 [58], using R studio version 1.0.136 [59] and the ggplot2 package [60].

#### Gene based calibration curve

For the target amplicons of G. *lapillus*, a DNA fragment spanning the target sequence, the reverse and forward primer sites and an extra 50bp on either end was synthesised called gBlocks ®; by Integrated DNA Technologies (IDT, IA, USA). Lyophilized gBlocks ®; was re-suspended in 1x TE (Tris 1M, EDTA 0.5 *Λ* 0 pH8) to a concentration of 1 ng/μl. The copy number of gene fragment was then calculated as 2.88×10^10^ for *G. lapillus.* The stock solution was serially diluted (10-fold) and dilutions between 10^3^ and 10^8^ were amplified by qPCR (on StepOnePlus System by Applied Biosystems (Thermo Fisher Scientific, Waltham, MA, USA) in triplicate.

#### Cell based calibration curve

Two strains of *G. lapillus* (HG4 and HG7) were used to construct cell based standard curves. Cells were counted under a Nikon Eclipse TS100 (Australia) microscope using a Sedgwick Rafter counting chamber. DNA was extracted with the FastDNA spin kit for soil by MP Biomedicals (CA, USA), as per the manufacturer’s instructions. The gDNA extracts were 10-fold serially diluted. Dilutions ranging from 3880 to 0.04 cells and 5328 to 0.05 for HG4 and HG7 respectively. Samples were amplified via qPCR (on StepOnePlus System by Applied Biosystems (Thermo Fisher Scientific, Waltham, MA, USA) in triplicate.

### Determination of gene copies per cell for *G. lapillus*

To determine the mean SSU rDNA copies per cell, the dilution series with a known cell count (3880 to 0.04 cells and 5328 to 0.05 for HG4 and HG7 respectively, see section *Cell based calibration curve)* were used as input for calculation.

The slope of the linear regression of SSU copies was used to determine copy number by correlating the qPCR detection of the gene based calibration curves and cell numbers. This slope of the linear regression was then used to determine the gene copy number per cell [61].

### Screening environmental samples for *G. lapillus*

Around Heron Reef (Fig. 2) 25 sites (within 1km of the shore) were sampled in October 2015, in spatial replicates (A, B, C) within a 2m radius. Representatives of three genera of macroalgae that commonly grow on this reef, *Chnoospora* spp, *Padina* sp. and *Saragassum* sp., were sampled for the presence of epiphytic *Gambierdiscus* spp. For each sample, about 200 g of macroalgae was collected from approximately 1 m deep water at low tide and briefly placed in plastic bags containing 200 to 300 ml of ambient seawater. They were shaken vigorously for 5 min to detach the epiphytic dinoflagellates from the macroalgae. This seawater was passed through > 120 m mesh filter to remove any remaining larger fauna and debris. The collected seawater was centrifuged at 1000 rpm. The supernatant was discarded and the pellet was dissolved in 10 ml RNAlater (Ambion, Austin, TX, USA) for preservation and stored at 4° C. Samples were screened in triplicate for both G. *lapillus* on a StepOnePlus System by Applied Biosystems (Thermo Fisher Scientific, Waltham, MA, USA).

**Figure 2:**
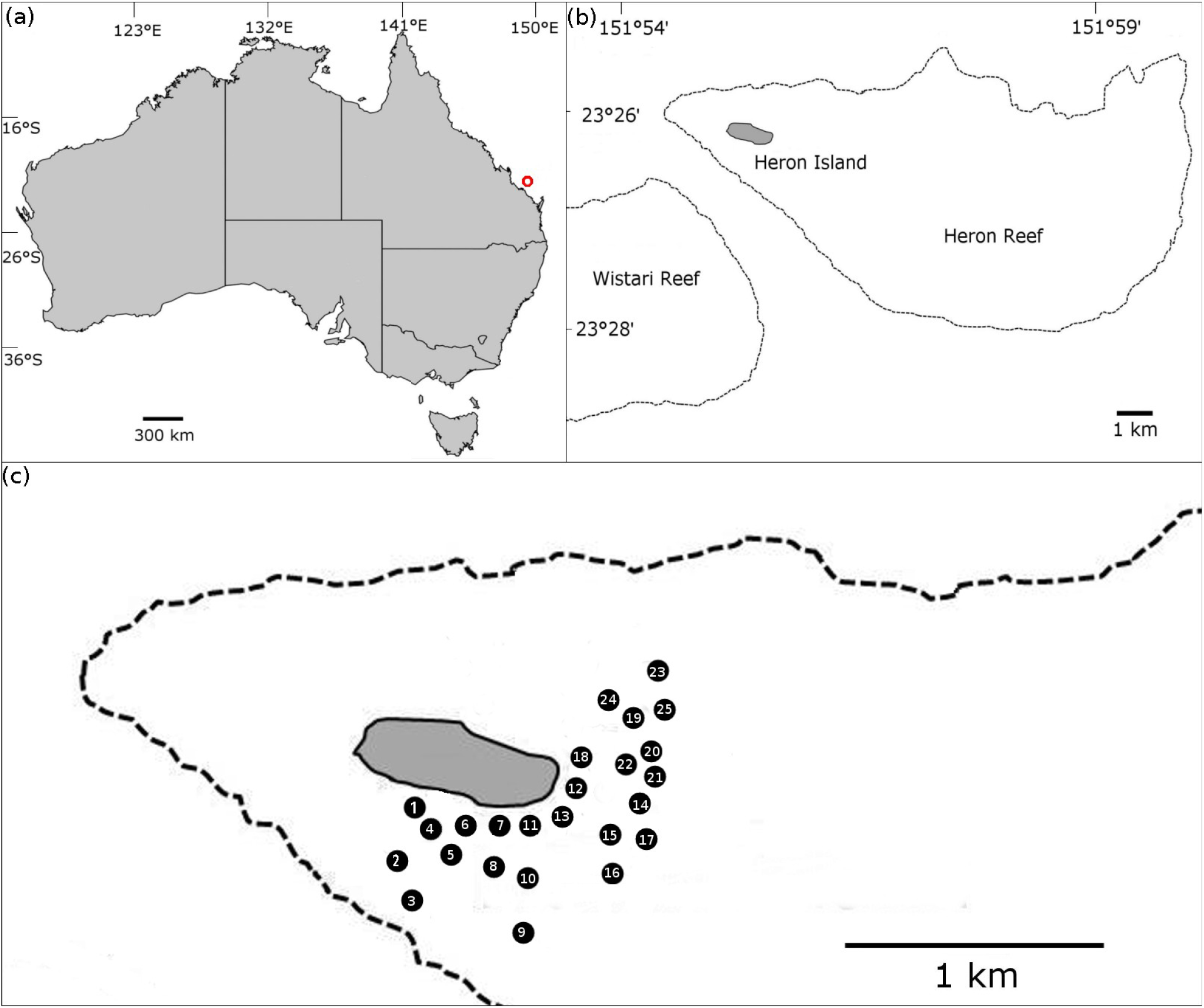
(A) Map of Australia, with the position of Heron Island (red circle); (B) Heron Island including surrounding reefs; (C) Approximate location of sampling sites around Heron Island. Map adapted from Kretzschmar et al. (2017) [19] and edited in the GNU Image Manipulation Program 2.8 (http://gimp.org)

## Results

### Evaluation of primer specificity

The qGlapSSU2F – qGlapSSU2R primer pair (Table 4) produced an amplified product of 138 bp for all five strains of G. *lapillus*, while no amplification was observed for genetically closely related species G. *belizeanus*, G. *cheloniae*, G. *pacificus* and *G. scabrosus.* Other species of *Gambierdiscus* from different clades, G. *australes*, G. *carpenteri*, G. *polynesiensis* and G. cf. *silvae* (Table 5) were also not amplified using this primer set [11, 19].

**Table 5:** Cross-reactivity of the qPCR primer set based on presence absence of PCR product visualised in agarose gel.

| Template | Strain name | gDNA gel band | GlapSSU2F-GlapSSU2R |
| --- | --- | --- | --- |
| *G. australes* | CCMP1650 | + | - |
|  | CG61 | + | - |
| *G. belizeanus* | CCMP401 | + | - |
| *G. carpenteri* | UTSMER9A3 | + | - |
| *G. cheloniae* | CAWD232 | + | - |
| *G. lapillus* | HG1 | + | + |
|  | HG4 | + | + |
|  | HG6 | + | + |
|  | HG7 | + | + |
|  | HG26 | + | + |
| *G. pacificus* | CAWD149 | + | - |
| *G. polynesiensis* | CG14 | + | - |
|  | CG15 | + | - |
| *G. scabrosus* | KW070922_1 | + | - |
| *G. cf. silvae* | HG5 | + | - |

### Evaluation of primer sensitivity

The cell-based standard curves for G. *lapillus* (HG4 and HG7, Fig. 3a) showed high linearity with R^2^ approaching 1.00. The slope for the Ct vs. log _10_ cell for HG4 was −3.4, which corresponds to an efficiency 96.8 %; and −3.51, which corresponds to an efficiency of 92.7 % for HG7 (Fig. 3). The linear detection for both G. *lapillus* isolates covered six orders of magnitude. The lowest number of cells detected were 0.04 and 0.05 cells for HG4 and HG7 respectively (Fig. 3a).

**Figure 3:**
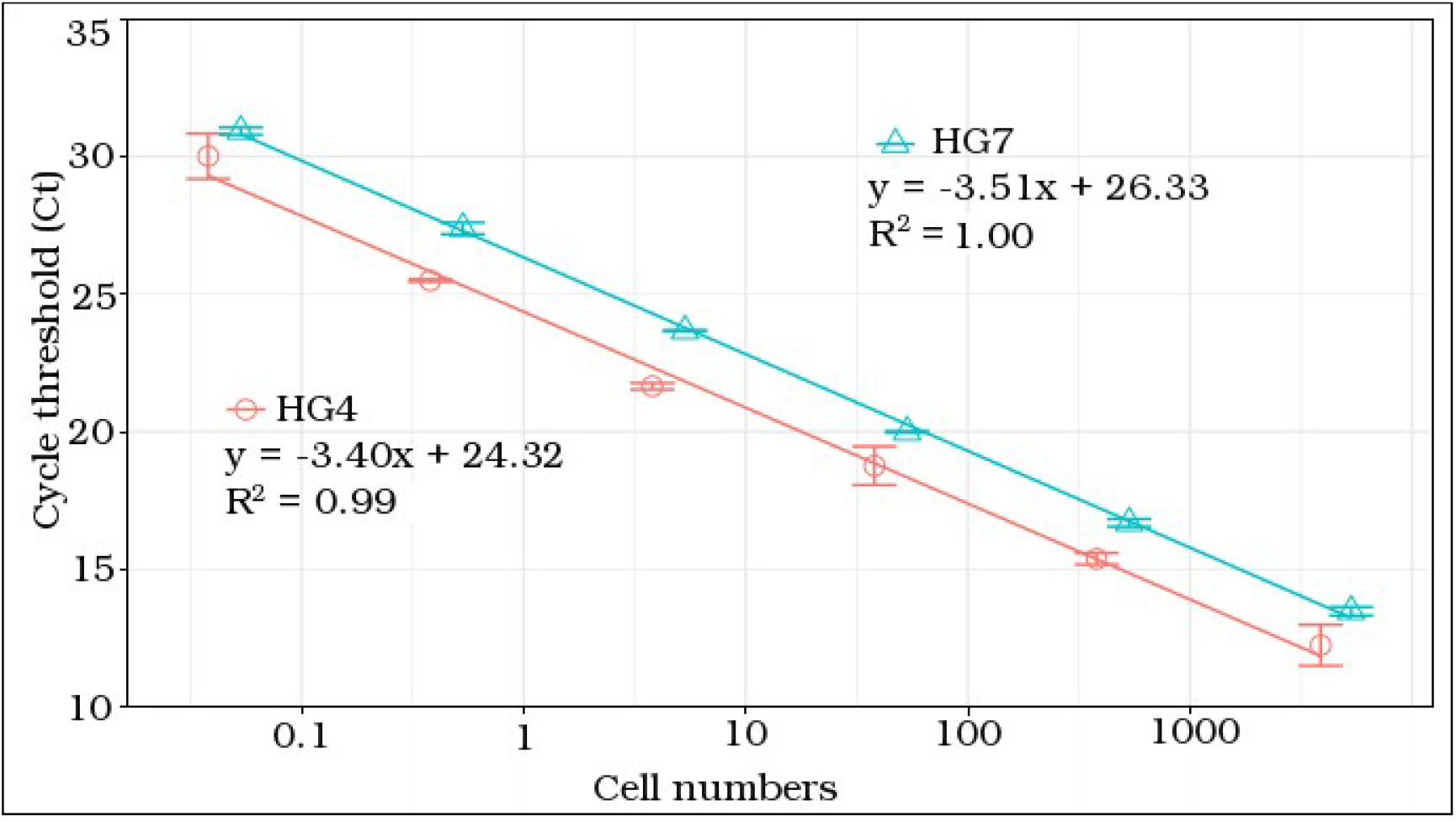
qPCR cell based standard curves of G. *lapillus* strains HG4 (circle) and HG7 (triangle). Error bars represent the deviation of technical replicates during reactions.

The gene based (gBlocks) standard curve for *G. lapillus* covered linear detection over 7 orders of magnitude, with a slope of −3.42, and a PCR efficiency of 96 % (Fig. 4). The detection limit tested was less than 10^5^ gene copy numbers. The Ct for the lowest gene copy number tested was less than 25, so it is likely that the sensitivity is lower than 10^5^ gene copy numbers (Fig. 4).

**Figure 4:**
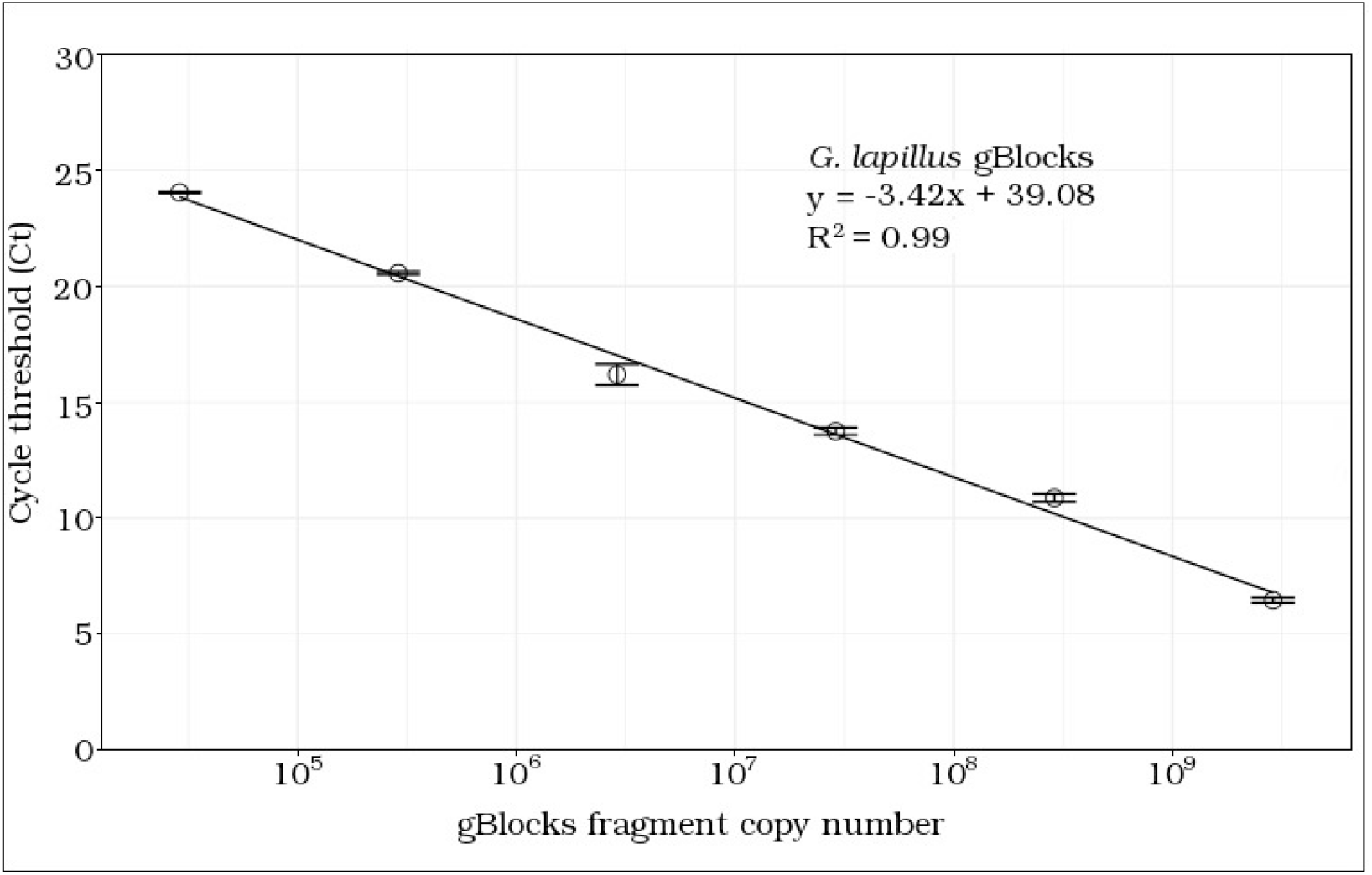
qPCR gene based standard curves of G. *lapillus.* Error bars represent the deviation of technical replicates during reactions.

### Quantification of SSU rDNA copy number per cell of *G. lapillus*

The detectable SSU copies for *G.lapillus* were 2.24 x 10^4^ and 5.85 x 10^3^ copies per cell for HG4 and HG7 respectively.

### Screening environmental samples for *G. lapillus* abundance

To evaluate the adequacy of the G. *lapillus* qPCR assay for environmental screening, the assay was applied to environmental community DNA extracts collected around Heron Island (Fig. 2). A relatively low cell abundance was detectable for G. *lapillus.* Ct values for G. *lapillus* detection in environmental samples were calibrated to the HG7 standard curve and calculated as cells.g^-1^ wet weight macroalgae (Table 6). *G. lapillus* was detected across 24 of the 25 sampling sites. At sites at which *G. lapillus* was present, it showed a patchy distribution, being present at two of the three spatial replicates in the majority of samples (17 of 25 sample sites), followed by all three spatial replicates testing positive (6 out of 25 sites) and at one site only one of the spatial replicates was positive (Fig. 5).

**Figure 5:**
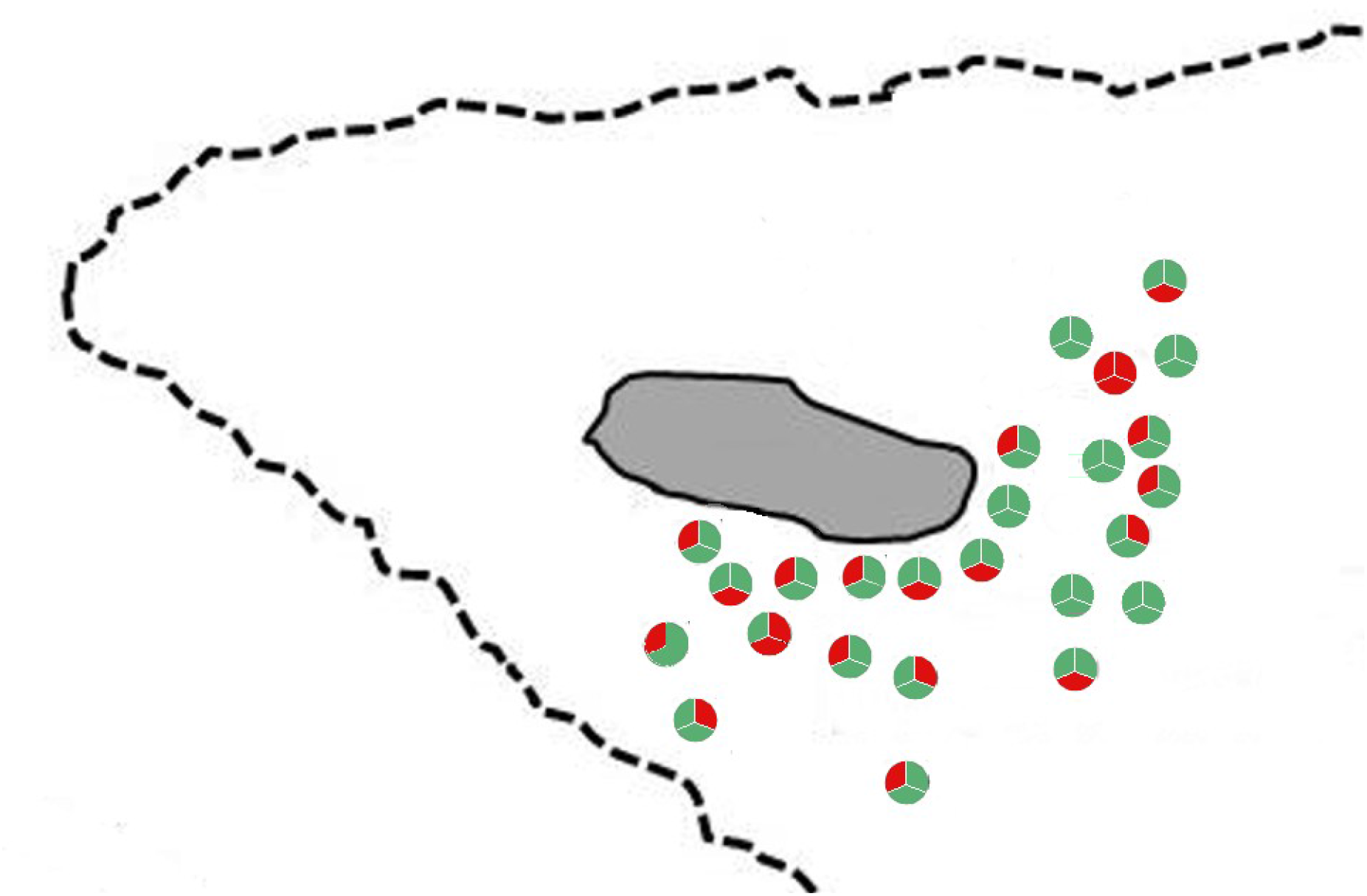
G. *lapillus* presence at the macroalgal sampling sites around Heron Island. The spatial replicates for each site are set up as shown in (A); the sites in (B) linked to numbering in Fig. 2 where positive (green) and negative (red) as per Table 6.

**Table 6:**
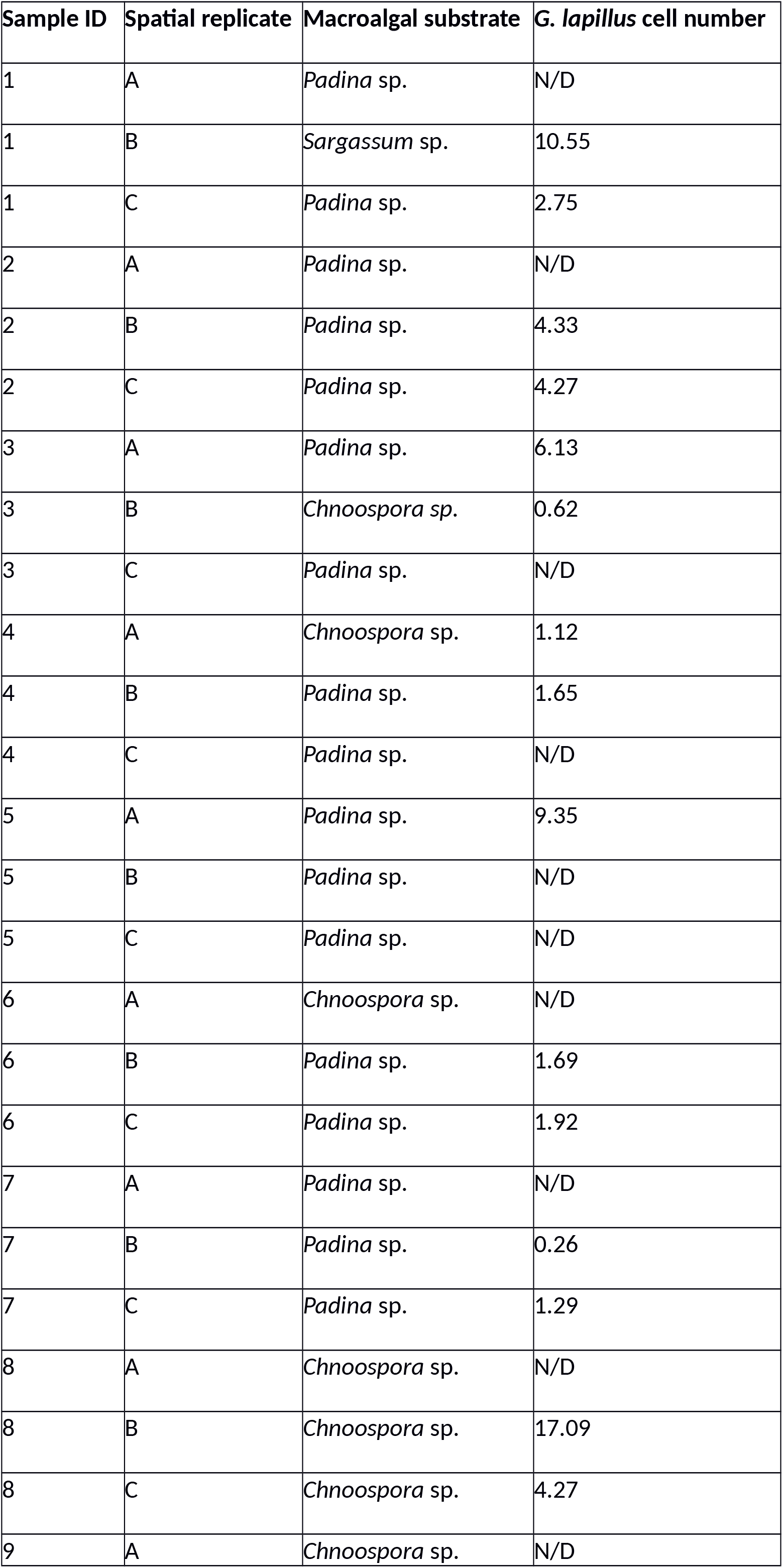
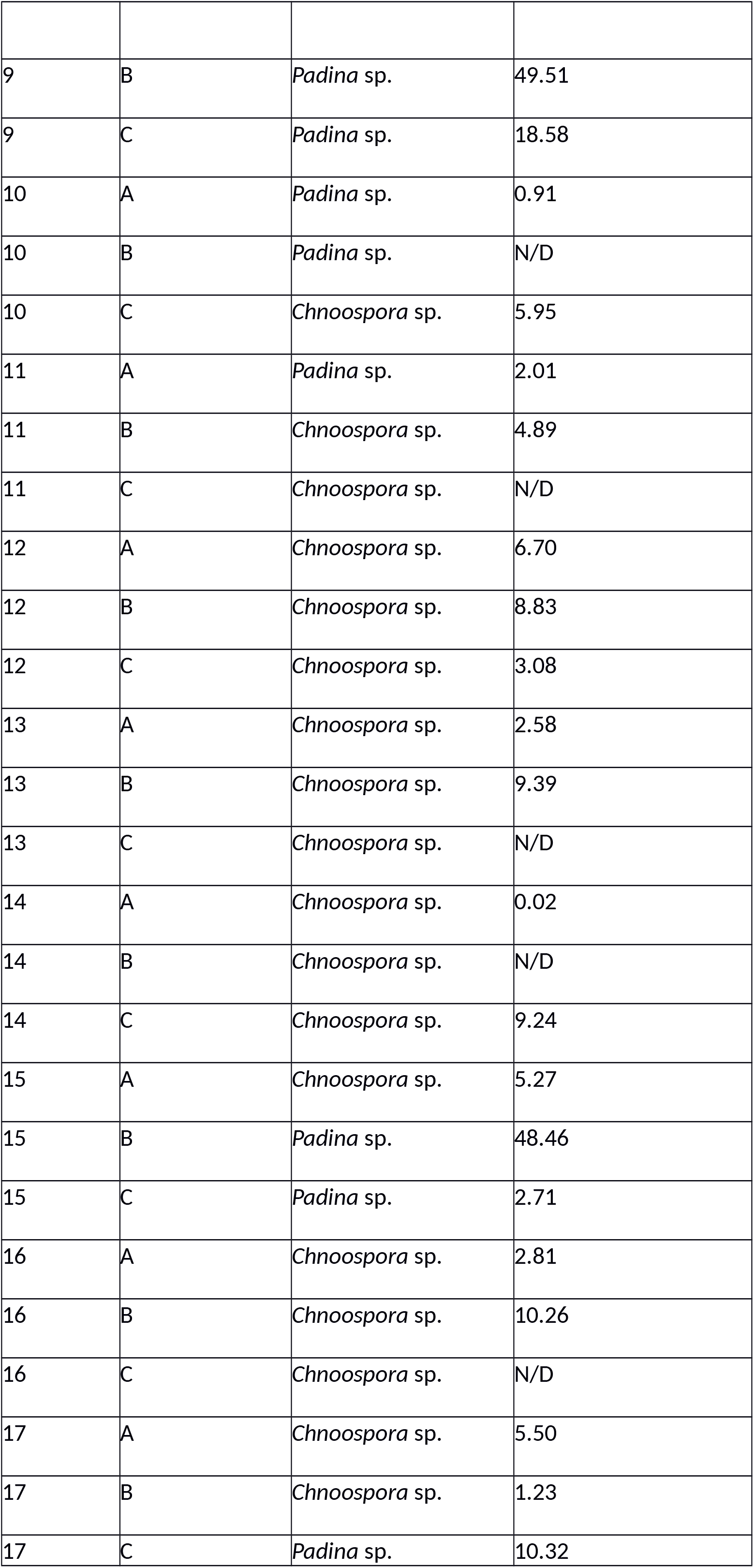
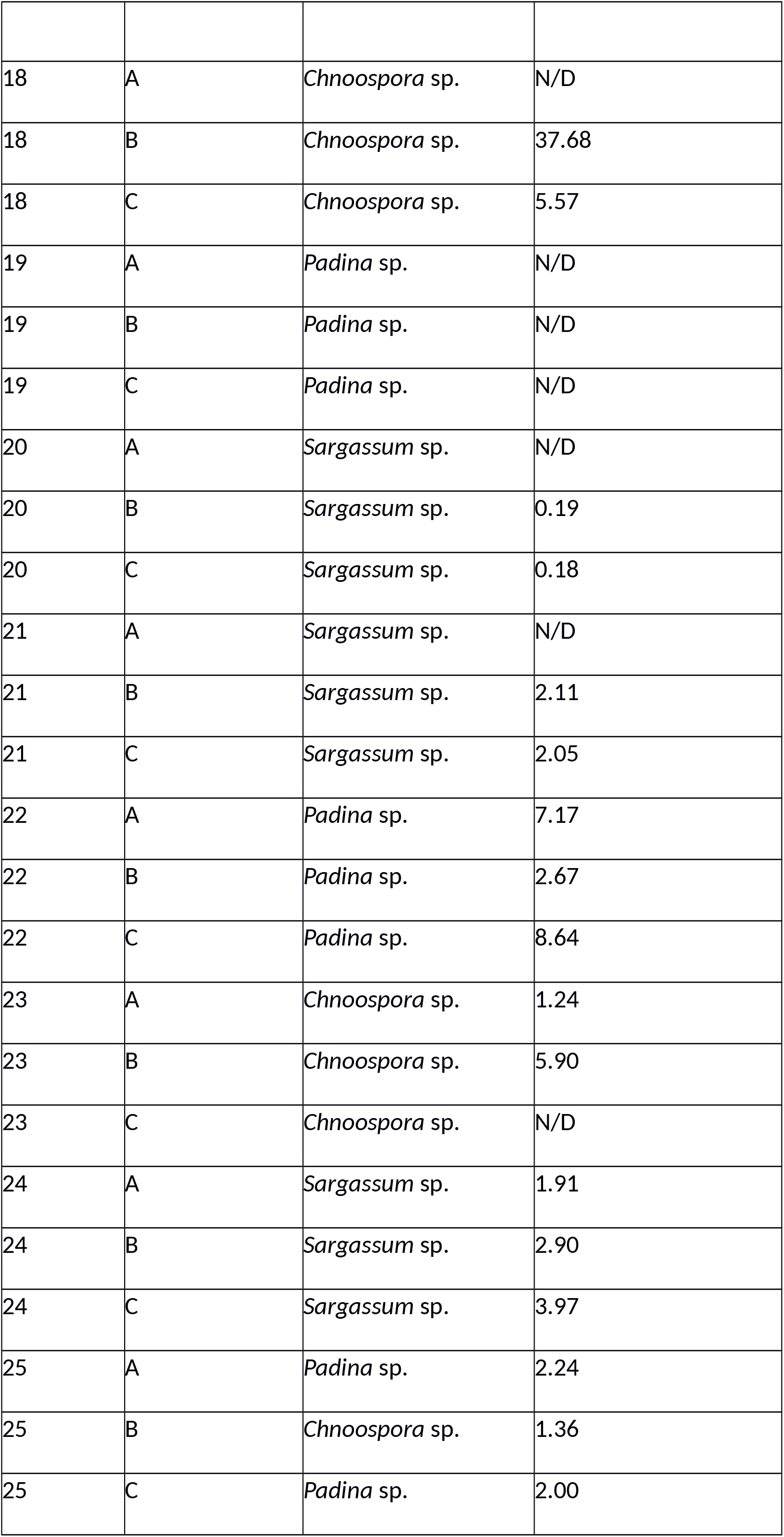
Screening of macroalgal samples for G. *lapillus* and cell density estimates via qPCR. Cell numbers were modeled on the type strain HG7. N/D denotes not detected; N/A denotes not attempted due to loss of sample.

| Sample ID | Spatial replicate | Macroalgal substrate | *G. lapillus* cell number |
| --- | --- | --- | --- |
| 1 | A | *Padina* sp. | N/D |
| 1 | B | *Sargassum* sp. | 10.55 |
| 1 | C | *Padina* sp. | 2.75 |
| 2 | A | *Padina* sp. | N/D |
| 2 | B | *Padina* sp. | 4.33 |
| 2 | C | *Padina* sp. | 4.27 |
| 3 | A | *Padina* sp. | 6.13 |
| 3 | B | *Chnoospora* sp. | 0.62 |
| 3 | C | *Padina* sp. | N/D |
| 4 | A | *Chnoospora* sp. | 1.12 |
| 4 | B | *Padina* sp. | 1.65 |
| 4 | C | *Padina* sp. | N/D |
| 5 | A | *Padina* sp. | 9.35 |
| 5 | B | *Padina* sp. | N/D |
| 5 | C | *Padina* sp. | N/D |
| 6 | A | *Chnoospora* sp. | N/D |
| 6 | B | *Padina* sp. | 1.69 |
| 6 | C | *Padina* sp. | 1.92 |
| 7 | A | *Padina* sp. | N/D |
| 7 | B | *Padina* sp. | 0.26 |
| 7 | C | *Padina* sp. | 1.29 |
| 8 | A | *Chnoospora* sp. | N/D |
| 8 | B | *Chnoospora* sp. | 17.09 |
| 8 | C | *Chnoospora* sp. | 4.27 |
| 9 | A | *Chnoospora* sp. | N/D |
| 9 | B | *Padina* sp. | 49.51 |
| 9 | C | *Padina* sp. | 18.58 |
| 10 | A | *Padina* sp. | 0.91 |
| 10 | B | *Padina* sp. | N/D |
| 10 | C | *Chnoospora* sp. | 5.95 |
| 11 | A | *Padina* sp. | 2.01 |
| 11 | B | *Chnoospora* sp. | 4.89 |
| 11 | C | *Chnoospora* sp. | N/D |
| 12 | A | *Chnoospora* sp. | 6.70 |
| 12 | B | *Chnoospora* sp. | 8.83 |
| 12 | C | *Chnoospora* sp. | 3.08 |
| 13 | A | *Chnoospora* sp. | 2.58 |
| 13 | B | *Chnoospora* sp. | 9.39 |
| 13 | C | *Chnoospora* sp. | N/D |
| 14 | A | *Chnoospora* sp. | 0.02 |
| 14 | B | *Chnoospora* sp. | N/D |
| 14 | C | *Chnoospora* sp. | 9.24 |
| 15 | A | *Chnoospora* sp. | 5.27 |
| 15 | B | *Padina* sp. | 48.46 |
| 15 | C | *Padina* sp. | 2.71 |
| 16 | A | *Chnoospora* sp. | 2.81 |
| 16 | B | *Chnoospora* sp. | 10.26 |
| 16 | C | *Chnoospora* sp. | N/D |
| 17 | A | *Chnoospora* sp. | 5.50 |
| 17 | B | *Chnoospora* sp. | 1.23 |
| 17 | C | *Padina* sp. | 10.32 |
| 18 | A | *Chnoospora* sp. | N/D |
| 18 | B | *Chnoospora* sp. | 37.68 |
| 18 | C | *Chnoospora* sp. | 5.57 |
| 19 | A | *Padina* sp. | N/D |
| 19 | B | *Padina* sp. | N/D |
| 19 | C | *Padina* sp. | N/D |
| 20 | A | *Sargassum* sp. | N/D |
| 20 | B | *Sargassum* sp. | 0.19 |
| 20 | C | *Sargassum* sp. | 0.18 |
| 21 | A | *Sargassum* sp. | N/D |
| 21 | B | *Sargassum* sp. | 2.11 |
| 21 | C | *Sargassum* sp. | 2.05 |
| 22 | A | *Padina* sp. | 7.17 |
| 22 | B | *Padina* sp. | 2.67 |
| 22 | C | *Padina* sp. | 8.64 |
| 23 | A | *Chnoospora* sp. | 1.24 |
| 23 | B | *Chnoospora* sp. | 5.90 |
| 23 | C | *Chnoospora* sp. | N/D |
| 24 | A | *Sargassum* sp. | 1.91 |
| 24 | B | *Sargassum* sp. | 2.90 |
| 24 | C | *Sargassum* sp. | 3.97 |
| 25 | A | *Padina* sp. | 2.24 |
| 25 | B | *Chnoospora* sp. | 1.36 |
| 25 | C | *Padina* sp. | 2.00 |

*G. lapillus* was detected at 71 out of the 75 spatial replicates, specifically at 24/32, 22/33 and 8/10 samples from *Chnoospora* sp., *Padina* sp. and *Saragassum* sp. as substrate respectively (Table 6). Patchiness was also found in the abundance as well as the distribution of G. *lapillus*, from 0.24 cells.g^-1^ wet weight macroalgae to 49.51 cells.g^-1^ wet weight macroalgae, with a mean of 5.84 cells.g^-1^ wet weight macroalgae. For example (4A – *Chnoospora* sp.) and (4B – *Padina* sp.) hosted comparable cell numbers (1.12 cells and 1.65 cells.g^-1^ wet weight algae respectively) while no G. *lapillus* cells were detected on (4C – *Padina* sp.). Only at one of 25 sampling sites, no G. *lapillus* presence was detected across all three spatial replicates (19A, B, C). At all other sites, the presence of G. *lapillus* varied between spatial replicates but did not significantly differ between macroalgal host or location (Fig. 6).

**Figure 6:**
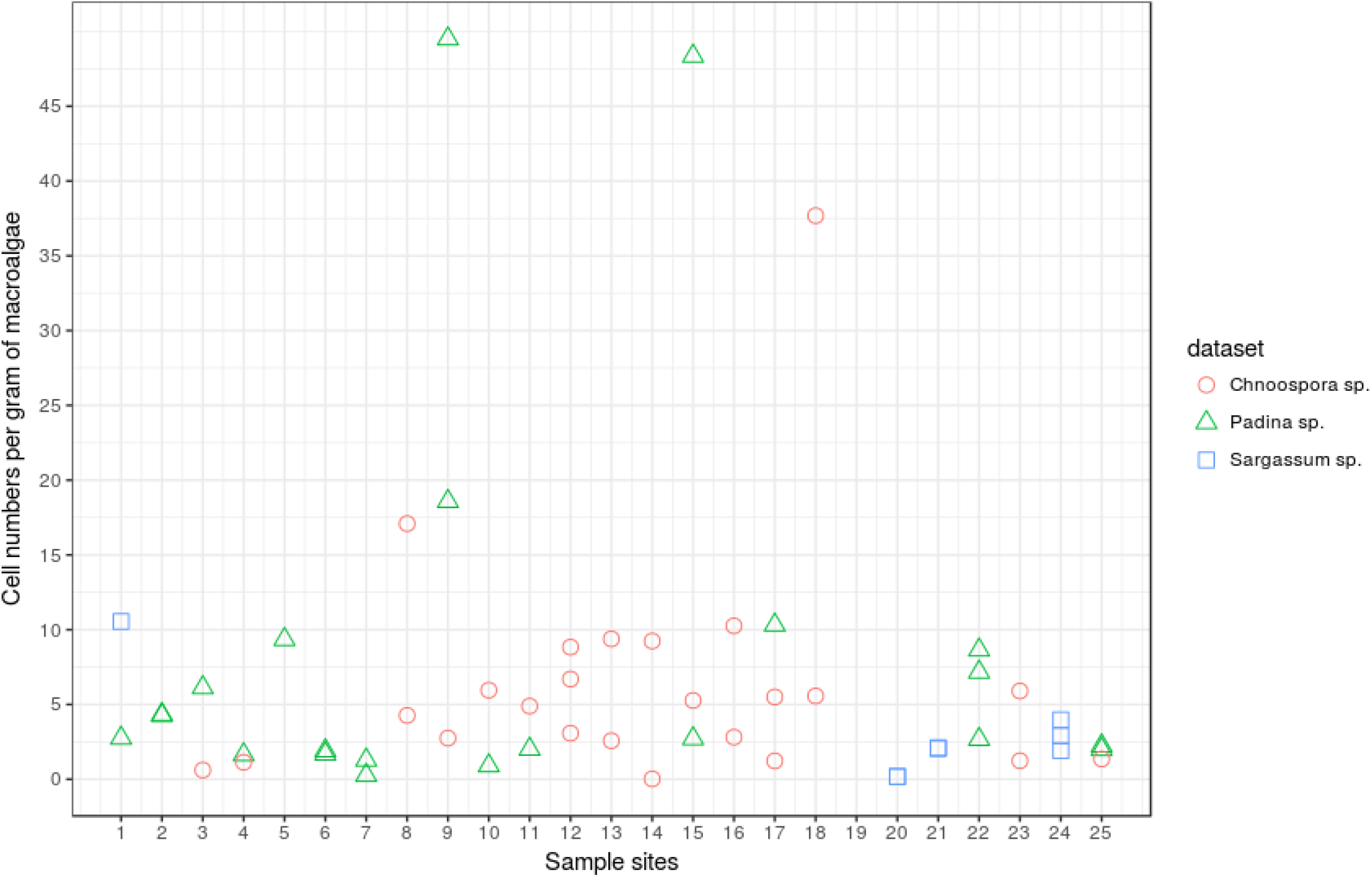
Detection of G. *lapillus* per spatial replicate at each macroalgal sampling site. Cell numbers were normalised to the HG7 standard curve (Fig. 3A). Figure also shows spatial replicates per macroalgal substrate where *Chnoospora* sp. samples are represented by circles, *Padina* sp. by squares and *Sargassum* by crosses (Table 6).

## Discussion

The aim of the study was to design and validate a species-specific qPCR assay to quantify *G. lapillus* a species that may produce CTX-like toxicity in the Australian GBR region. Species-specific qPCR primers with high specificity and sensitivity were developed and the SSU copy number for two strains were determined, and were found to differ from one another considerably, as one strain had more than four times the number of genomic rDNA copies. We also established that this primer set was effective in measuring the abundance and distribution of G. *lapillus* at the Heron Island reef. The cross-reactivity of primers designed in this study showed high specificity for both G. *lapillus* while not amplifying when tested against other *Gambierdiscus* spp. The species tested for cross-reactivity were chosen because they represented species that are genetically most similar to each target species for the SSU region (as per Fig. 2 in [19]). Standard curves were constructed for two strains of G. *lapillus* for which the primers showed high linearity and amplification efficiency (Fig. 3). Hence, this primer set is an accurate and reproducible molecular tool to enumerate the target species exclusively from environmental community DNA extracts. More importantly, this assay does not require the operator to rely on melt curves to identify species, or to have access to G. *lapillus* DNA extracts as a positive control. Due to the potential CTX production of G. *lapillus* [19, 48] the presence and distribution of this species is of interest in Australia where the causative organism(s) for CFP is yet to be established.

As CFP risk is linked to the abundance of *Gambierdiscus* species producing CTXs [32, 62], it was important to establish a quantitative assay for detection. We validated a synthetic gene fragment standard curve of the target region (gBlocks ®;) and compared this to cell standard curves to establish an ‘absolute’ qPCR assay [40, 63]. Further, we determined the copy SSU rDNA number for two strains of G. *lapillus* (HG4 and HG7). The copy number for G. *lapillus* (5,855.3 to 22,430.3 rDNA copies per cell) were comparable to the copy numbers determined by Vandersea et al. (2012), which ranged from 690 rDNA copies for G. *belizeanus* to 21,498 copies for G. *caribaeus.* In comparison, the cell copy numbers determined by Nishimura et al. (2016) ranged from 532,000 copies for G. *scabrosus* and 2,261,000 for *G.* sp. type 3. While the difference in rDNA copy numbers may be due to inter-species differences, or even intra-species as per the *G. lapillus* results, Nishimura et al. (2016) argue that the difference could be underestimation of rDNA copy numbers due to ‘ghost’ cells counted for total cell number which do not contribute to amplification [40, 63]. The difference in SSU rDNA copies between the two strains of G. *lapillus* isolated from the same region highlights the importance of carefully verifying qPCR assays based on rRNA genes using multiple local strains. A difference of this magnitude may lead to considerably different abundance estimates of G. *lapillus.* As the variation between the two strains tested is within the observed variation reported by Nishimura et al. (2016) from single cell qPCR experiments for rDNA copy number elucidation, the difference reported here is likely representative of biological intra-strain variation rather than methodological artifacts. A 5-fold difference in toxicity between the same HG4 and HG7 strains for G. *lapillus* was also reported by Kretzschmar et al. (2017), and there was a noticeable difference in growth rate between the two strains observed (but not quantified) in this study. The mounting evidence of intra-strain variability in toxicity, detectable rDNA copy numbers and potentially growth rate could have severe implications for qPCR based cell enumeration of environmental samples when attempting to extrapolate CFP risk and requires further investigation.

The qPCR assay was successfully tested on environmental DNA extracts from around Heron Island, and gave some insight into G. *lapillus* distribution and abundance. The qPCR assay detected G. *lapillus* at all of the sites tested (Fig. 5). Within the spatial replicates, the distribution of G. *lapillus* was patchy, as 24 of the 25 sites included at least one replicate with no G. *lapillus* present (Fig. 5). Patchiness in the distribution of *Gambierdiscus* species has previously been found in a study of 7 *Bryothamnion* macroalgae spaced 5 to 10 cm apart, in which 5 to 70 cells g-1 algae were found [64].

There was no significant difference in the presence/absence of G. *lapillus* cells observed as per the macroalgal host, *Padina* sp. or *Sargassum* sp.

Motile behaviour has been observed previously in the field at various time points [65, 66]. Parsons et al. (2011) reported *Gambierdiscus* sp. behaviour as facultative epiphytes during lab scale experiments, as cells showed attachment as well as motile stages over time in the presence of different macroalgae [67]. Taylor & Gustavson (1983) reported that *Gambierdiscus* cells were captured in plankton tows by de Silva in 1956 but reported as *Goniodoma* [64]. Motility could be a factor for the patchy distribution observed in the spatial replicates. Across spatial replicates where G. *lapillus* was detected, cell densities were consistent (Fig. 6). The average cell density of G. *lapillus* 5.84 cells.g^-1^ wet weight macroalgae, which is comparable to the cell densities recorded by Nishimura et al. (2016) in their environmental screening (Table 4 in [40]).

As many authors have pointed out (e.g. [13, 32, 67, 68, 69, 70, 71]), there are several difficulties in determining precise quantification of *Gambierdiscus* species on macroalgae in order to assess potential CFP risk. Due to the difference in habitable surface area between samples taken from structurally diverse macroalgae, including those sampled in this study *(Chnoospora, Padina* sp. and *Sargassum* sp.), the potential habitable space is difficult to compare. Further, in order to assess CFP risk in a given area, the properties of the macroalgae with *Gambierdiscus* spp. epiphytes need to be considered. If the macroalgae is structurally or chemically defended against herbivory, any CTX produced by the epiphytes is unlikely to enter the food chain and cause CFP [70]. Due to the difficulty in quantifying *Gambierdiscus* spp. on a particular substrate, Tester et al. (2014) proposed have the use of an artificial substrate (commonly available black fibreglass screen of a known surface area) and a standardised sampling method [69].

## Conclusion

The qPCR assay developed in this study is an accurate molecular tools to detect and enumerate the presence of G. *lapillus* in environmental samples. The assay was shown to be highly sensitive and accurately detected 0.05 to over 4000 cells for G. *lapillus.* Although the toxin profile of G. *lapillus* has not been completely defined, it may produce uncharacterised CTXs congeners [19, 48] and therefore is a part of the ciguateric web in Australia. The assay was applied to spatial replicates from 25 sites around Heron Island on the GBR, which found that G. *lapillus* was commonly present, but had a patchy spatial distribution and abundance. The development and validation of a quantitative monitoring tool presented here for G. *lapillus* is in line with Element 1 of the Global Ciguatera Strategy [32].

## Acknowledgements

We are grateful to Dr. Adachi and Dr. Nishimura for supplying G. *scabrosus*, Michaela Larsson for supplying *G. carpenteri* DNA as well DNA as Dr. Kirsty Smith and Dr. Lesley Rhodes for supplying G. *cheloniae* DNA. DNA from all three species was used for cross reactivity assessment.

A.L.K. was supported by a UTS doctoral scholarship funded by the University of Technology Sydney. S.M and G.Ks involvement was supported by the Australian Research Council grant FT120100704.

## Conflict of interest

The authors report no conflict of interest in conducting this study.

## Author contribution

The project and experiments were conceived by A.L.K and S.M. The environmental samples were collected by G.K. DNA from environmental samples was extracted by A.L.K., A.V. and G.K. The G. *lapillus* assay was designed and tested by A.L.K. The manuscript was drafted by A.L.K. and revised by all authors.

